# The Activation of the Transcription Factor NRF2 in Epithelial Cells Lining the Kidney Cysts in Tuberous Sclerosis Complex (TSC): The Interplay of Fumarate Hydratase 1, KEAP1, and NRF2 in Kidney Cystogenesis

**DOI:** 10.64898/2026.09.22.753536

**Authors:** Sharon Barone, Kamyar Zahedi, Manoocher Soleimani

## Abstract

**Background:** Tuberous Sclerosis Complex (TSC) is caused by inactivating mutations in either the *TSC1* or *TSC2* genes, leading to activation of the mammalian target of rapamycin complex 1 (mTORC1) and unhindered cell growth and proliferation. The epithelium of TSC renal cysts in both mice and humans is composed of proliferating A-intercalated (A-IC) cells. The exact molecular mechanism of kidney cystogenesis in TSC remains speculative.

**Hypothesis:** Superfluous cell proliferation driven by mTORC1 activation increases metabolic demand, causing oxidative stress and excess reactive oxygen species. If unchecked, this overwhelms antioxidant defenses and causes cell death. In TSC kidney cystogenesis, Nuclear Factor Erythroid 2-Related Factor 2 (NFE2L2), also known as NRF2, serves as the “master regulator” of antioxidant and anti-inflammatory responses, enabling cells to survive and proliferate by clearing the toxic environment and supplying nutrients and fuel.

**Results:** RNA-seq and proteomics, along with western blot analysis, showed robust downregulation of Fumarate Hydrase 1 (FH1) and upregulation of NRF2, STAT3, and HIF1α in kidneys from TSC mice with moderate or heavy cyst burden. Confocal microscopy and immunohistochemical staining on kidney sections, and/or western blot studies on nuclear and cytoplasmic fractions, showed nuclear localization of NRF2, STAT3, and HIF1α. In cyst-lining cells in TSC mouse models, FH1 downregulation was associated with inactivating succination of KEAP1 in immunoprecipitation experiments, promoting NRF2 nuclear localization in A-IC cells lining the cysts. NRF2 nuclear localization was associated with ectopic induction of the glutamine transporter SLC38A3 (SNAT3) on the basolateral membrane and activation of the NH_3_/NH_4_^+^ transporters RHCG and RHBG in A-IC cells lining the cysts. Twenty-four h urine NH_3_/NH_4_^+^ excretion rates increased significantly in *Tsc1* KO vs. WT mice.

The activation of NRF2, STAT3, and HIF1α can drive metabolic reprogramming and activate survival genes in proliferating cells. Together with SLC38A3 induction and upregulation of other glutamine and NH_3_/NH_4_ ^+^ transporters, these factors activate glutaminolysis and aerobic glycolysis, supplying nutrients to proliferating cystic epithelial cells and supporting cyst expansion in TSC. Consistent with this central role for glutaminolysis in kidney cystic epithelium and TSC cystogenesis, we find a significant reduction in kidney cyst burden in *Tsc1* KO mice on a glutamine-free diet.

**Conclusions:** NRF2 plays a critical role in antioxidant defense. Along with STAT3 and HIF1α, NRF2 is a key player in metabolic reprogramming through glutaminolysis, which supplies nutrients to proliferating cystic epithelial cells and supports cyst expansion in TSC. These findings suggest that inhibiting or inactivating NRF2, alone or in combination with HIF1α or STAT3, may represent a potential treatment strategy for kidney lesions in TSC

## Introduction

Tuberous sclerosis complex (TSC) is an autosomal dominant disease caused by inactivating mutations in the TSC1 or TSC2 genes, which encode Hamartin (TSC1) or Tuberin (TSC2), respectively (1–5). More than 2 million people are estimated to be living with TSC worldwide (2, 6). TSC affects multiple organs, including the kidney, lung, heart, and brain. In the kidney, TSC can present with cysts and angiomyolipomas, which eventually lead to kidney failure, thus requiring dialysis and/or a kidney transplant (3, 7, 8, 9). Despite our understanding of the genetic basis of TSC, the factors that promote renal lesion development (cysts and angiomyolipomas) remain poorly understood. Currently, rapalogs such as everolimus are the most effective treatments for TSC (3), but their efficacy is limited by serious side effects (10, 11), including recurrence of kidney lesions upon discontinuation (3, 9).

Mice with kidney principal cell (PC)-specific inactivation of *Tsc1* or *Tsc2* or pericyte-specific ablation of *Tsc1* genes develop numerous kidney cysts, which are primarily comprised of proliferating A-intercalated (A-IC) cells (12–17). The preponderance of A-IC cells in kidney cyst epithelium has also been documented in TSC patients (12, 13). In contrast, kidney cysts in Autosomal Dominant Polycystic Kidney Disease (ADPKD) are predominantly composed of non-A-IC cells (12, 18). Recent studies indicate that TSC kidney cystogenesis is promoted by enhanced expression of factors that regulate A-IC cell development, including Forkhead box transcription factors (FOXI1) and the receptor tyrosine kinase (c-KIT) (12, 19, 20).

Mechanistic Target of Rapamycin Complex 1 (mTORC1)-mediated cell proliferation, which is associated with increased metabolic demand, leads to oxidative stress and the generation of excess reactive oxygen species (ROS) (21–23). These changes initiate oxidative injury, and if not mitigated, lead to cell death. The transcription factor **nuclear factor erythroid-derived 2-like 2 (NFE2L2)**, also known as **NRF2**, is the primary sensor of oxidative stress and a “master regulator” of cellular antioxidant and anti-inflammatory responses (23–25), protecting cells under metabolic duress (26).

The purpose of these studies was to examine the expression and activation of NRF2 and its associated pathways and molecules in cystic epithelial cells in various *Tsc1* or *Tsc2* KO mice with mild, moderate, or heavy cyst burdens. Our results demonstrate that the transcription factor NRF2, as the primary sensor of oxidative stress and a “master regulator” of cellular antioxidant and anti-inflammatory response, is robustly activated and translocated to the nucleus in A-IC cells lining the cysts in *Tsc1* or *Tsc2* KO mice with moderate or heavy cyst burden. We further show the activation and nuclear localization of Signal Transducer and Activator of Transcription 3 (STAT3) and Hypoxia-Inducible Factor 1-alpha (HIF1α) in cells lining the cysts in TSC mouse models. These results further indicate activation of metabolic reprogramming (glutaminolysis and aerobic glycolysis) in cyst-lining cells in TSC. The significance of the results will be discussed.

## Results

### TSC mouse models

For our experiments, we used TSC mouse models with mild cyst burden (*Tsc1/Aqp2* Cre, 28 days old, moderate cyst burden (*Tsc1/Aqp2* Cre, 45 days old), or heavy cyst burden (*Tsc1/Car2/Aqp2 Cre* dKO, 110 days old) (12–17, 19). *Tsc1/Aqp2* Cre mice invariably die before 55 days of age and show a moderate kidney cyst burden at 45 days of age (12, 13). Immunohistochemical (IHC) staining experiments were performed to examine the expression of molecules identified by RNA-seq studies in these TSC mouse models, as well as *Tsc2/Aqp 2* Cre (135 days old) mice that have significantly longer lifespans (**Table 1**).

**Table 1.** TSC mouse models utilized in experiments. The TSC mouse type (Tsc1 or Tsc2) and official nomenclature is shown.

| Mouse Strains | Mouse Nomenclature |
| --- | --- |
| WT | C57BL6/J |
| ***Tsc1* Mice** |  |
| *Tsc1* KO | *Tsc1*<sup>tm1Djk</sup>/J **x** B6.Cg-Tg(Aqp2-cre)1Dek/J |
| *Tsc1*/Car2 dKO | *Tsc1*<sup>tm1Djk</sup>/J **x** B6.Cg-Tg(Aqp2-cre)1Dek/J **x** B6.D2-Ca2n/J |
| ***Tsc2* Mice** |  |
| *Tsc2* KO | *Tsc2*<sup>tm1.1Mjg</sup>/J **x** B6.Cg-Tg(Aqp2-cre)1Dek/J |

### RNA-Seq analysis, proteomics, and expression studies

RNA-seq data from kidneys of *Tsc1/Car2* dKO mice at 110 days of age, with heavy cyst burden, showed significant upregulation of NRF2 and downregulation of Fumarate Hydratase 1 (FH1). Immunohistochemical staining examined NRF2 expression in *Tsc1* KO and *Tsc2* KO mice (Table 1). **Figure 1A–C** shows robust nuclear localization of NRF2 in epithelial cells lining cysts in *Tsc1* KO (45 days old), *Tsc1/Car2* dKO (110 days old), and *Tsc2/Aqp2* Cre (125 days). For these IHC kidney sections, cyst burden is moderate or heavy. The kidneys of mice with mild cyst burden (*Tsc1* KO at 28 days of age) exhibited very few nuclear localizations (**Fig. 1D**). In contrast, the cystic epithelial cells in *Pkd1* KO mice showed scant nuclear NRF2 expression (**Fig. 1E**). In addition, our IHC studies showed NRF2 nuclear localization in kidney cyst tissues from an individual with TSC (**Fig. 1F**). Taken together, these results are consistent with NRF2 activation in kidney cyst epithelia in *Tsc1* and *Tsc2* KO mice with moderate or heavy cyst burden, but not in *Pkd1* KO mice.

**Figure 1.**
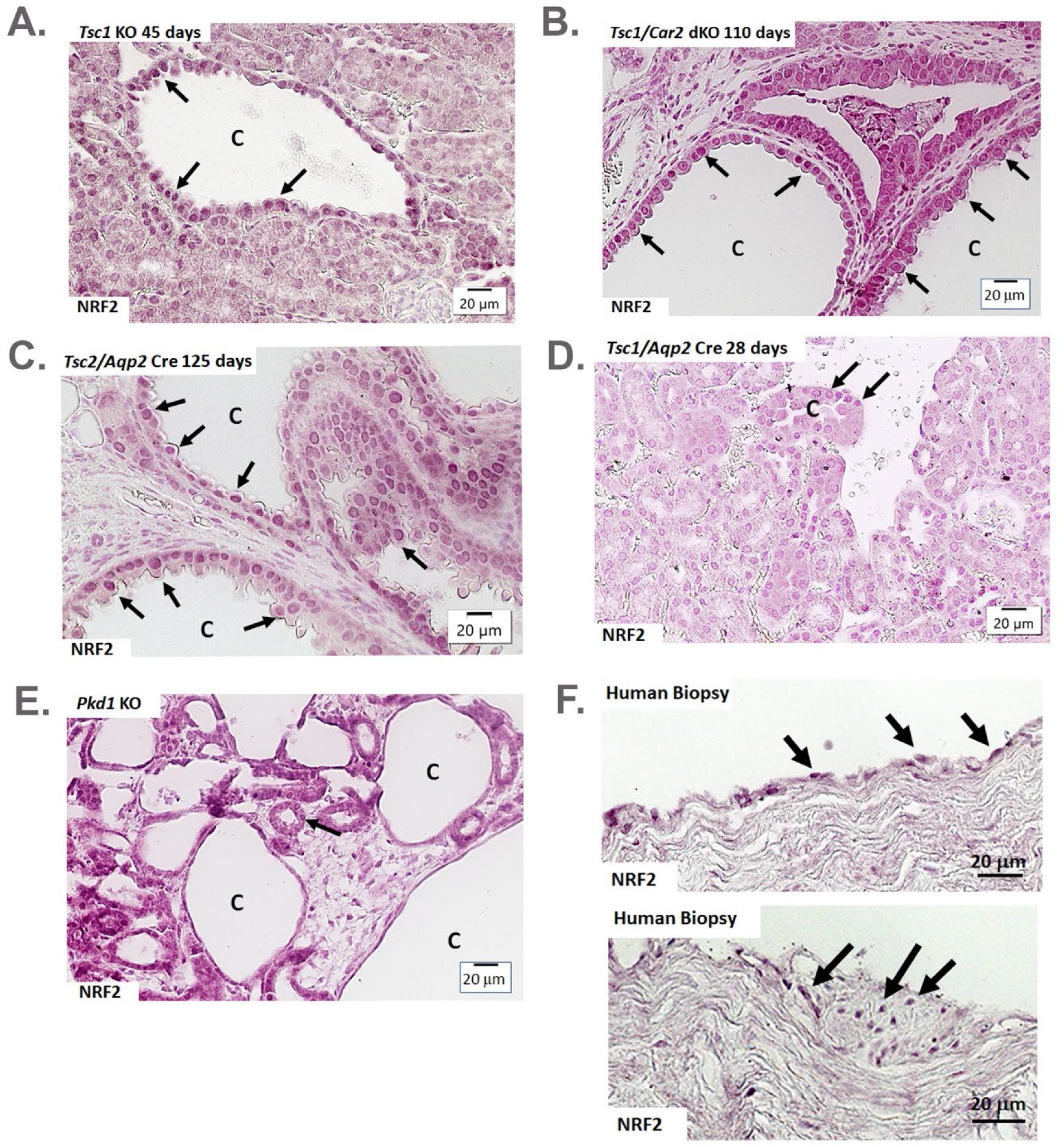
Nuclear expression of NRF2 in epithelial cells lining the cyst in TSC mouse models. Immunohistochemical labeling of NRF2 in **(A)** *Tsc1* KO (45 days of age), **(B)** *Tsc1/Car2* dKO (110 days of age), **(C)** *Tsc2* KO (125 days of age), **(D)** *Tsc1* KO (28 days of age), and **(E)** *Pkd1* KO mice. Black arrows point to nuclear localization of NRF2 in the epithelial cells lining the cysts. There is a lack of NRF2 present in the Pkd1 KO mouse. **E)** NRF2 expression in a cyst from an individual with TSC. Black arrows in the upper panel point to NRF2 nuclear localization in cystic cells facing the cyst lumen. Arrows in the lower panel point to nuclear localization in a section resembling angiomyolipoma. “C” represents cysts. Scale bar = 20μm.

### Confocal immunofluorescence microscopy

Using NRF2 and H^+^-ATPase antibodies, our studies demonstrated nuclear localization of NRF2 and apical expression of H^+^-ATPase in A-IC cells lining kidney cysts in *Tsc1* KO and *Tsc1/Car2* dKO **(Fig. 2A-B),** reaffirming that the cells lining TSC cysts are A-IC cells. Western blot analysis showed that NRF2 localized to the nuclear fractions prepared from cystic kidneys **(Fig. 2C).**

**Figure 2.**
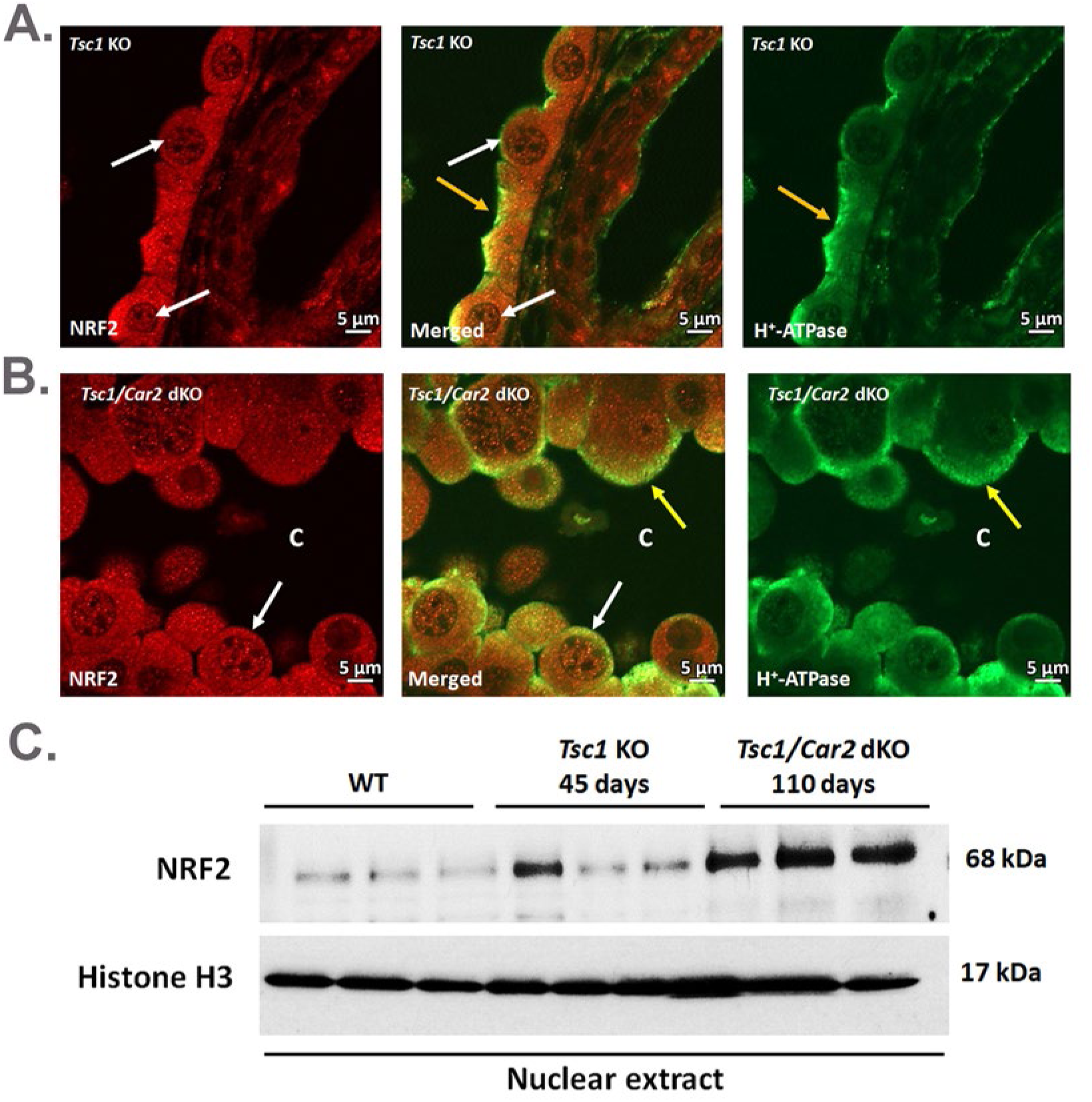
Localization and expression of NRF2 and H+-ATPase in the cystic epithelium of various TSC mouse models. Double-labeling of NRF2 (left panel; white arrows) and H^+^-ATPase (right panel; yellow arrows) lining the cysts in **(A)** Tsc1 KO and **(B)** *Tsc1/Car2* dKO mice. The middle panels illustrate the merged images. “C” represents cysts. Scale bar = 5μm. **C)** Western blot showing NRF2 expression in the nuclear extracts of *Tsc1* KO (45 days of age) and *Tsc1/Car2* dKO (110 days of age). Histone H3 displays equal loading of samples.

These results demonstrate NRF2 activation and nuclear localization in epithelial cells lining the cysts. Under normal conditions, Kelch-like ECH-associated protein 1 (KEAP1) regulates NRF2 and targets it for degradation (24–28). Our RNA-seq studies demonstrated significant downregulation of Fumarate Hydratase 1 (FH1) in mice with heavy cyst burden (*Tsc1/Car2* dKO mice at 110 days of age) vs. WT mice (p = 2.68x10^-10^), consistent with NRF2 activation. Western blot analysis confirmed FH1 downregulation in kidney lysate from mice with heavy cyst burden (*Tsc1/Car2* dKO mice at 110 days of age) (**Fig. 3A**). Proteomics studies (**Fig. 3B**) demonstrate FH1 downregulation in the kidneys of both *Tsc1* KO and *Tsc1/Car2* dKO mice, with robust downregulation in *Tsc1/Car2* dKO mice, consistent with the Western blot analysis in Fig. 3A.

**Figure 3.**
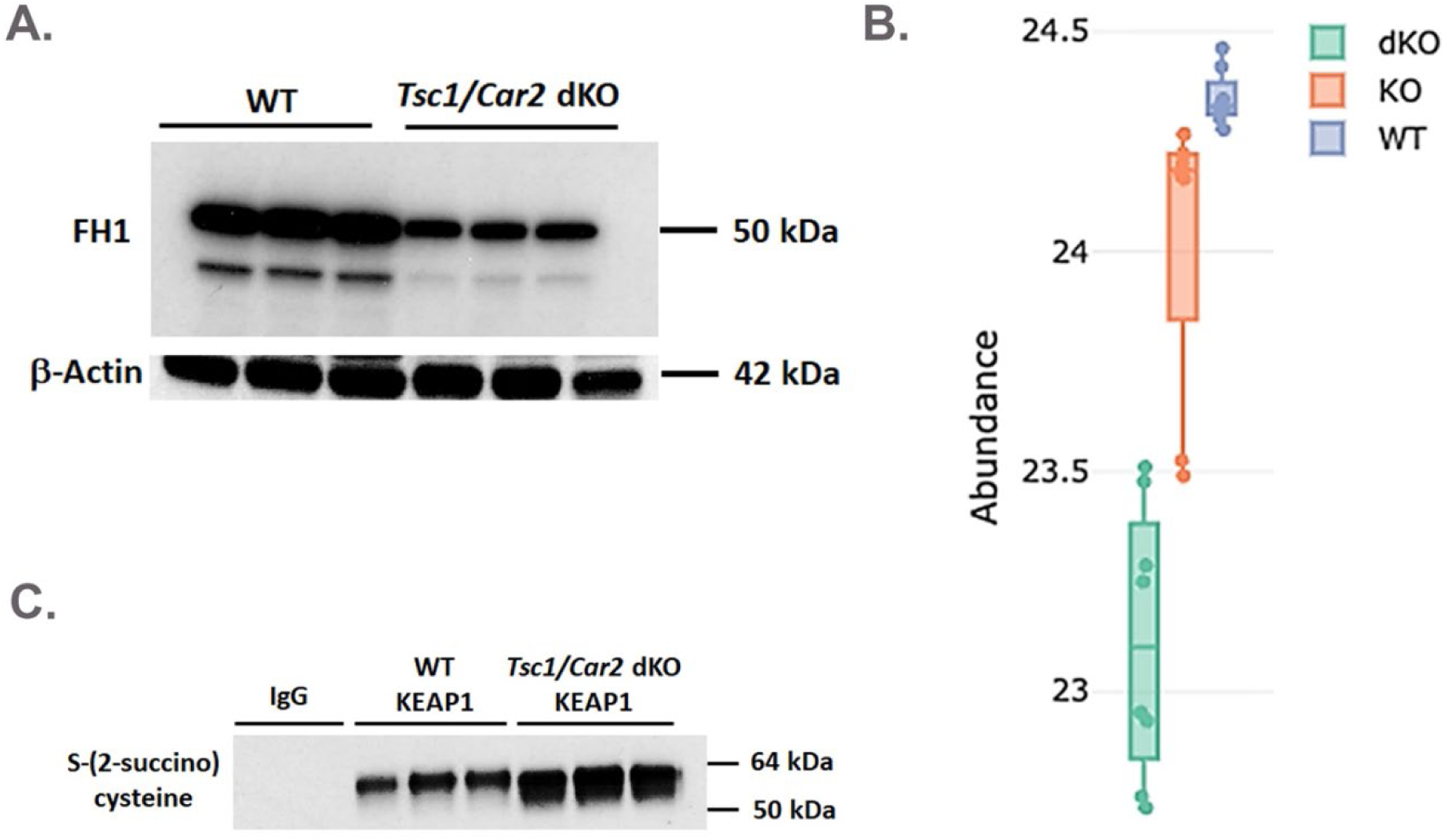
Protein expression of FH1 in TSC mouse models. **A)** Western blot demonstrating the decreased expression of FH1 in **Tsc1/Car2** dKO mice. **B)** Proteomic analysis of the kidneys of wildtype (WT), moderately cystic 45 days-old *Tsc1*KO (KO), and heavily cystic 110 days-old *Tsc1/Car2* dKO (dKO) mice. The expression of FH1 is significantly reduced in kidneys of KO and dKO mice vs. WT. **C)** KEAP1 was immunoprecipitated from kidneys of WT and *Tsc1/Car2*dKO mice using KEAP1 antibody and subsequently subjected to western blot analysis using an antibody to S-(2-succino) cysteine (2SC). The left 2 lanes are immunoprecipitation controls using pre-immune rabbit IgG1.

FH1 downregulation in tumors or disease states associated with mTORC1 hyperactivation results in accumulation of intracellular fumarate, which can drive succination of cysteine residues in KEAP1 (26–31). KEAP1 succination prevents it from binding to and mediating the proteasomal degradation of NRF2 (28–33). To examine the impact of FH1 downregulation on KEAP1 status, immunoprecipitation of KEAP1 was performed, and succination was analyzed by Western blot analysis. The results show strong KEAP1 succination (**Fig. 3C**). This loss of functional KEAP1 stabilizes NRF2, leading to its nuclear localization and binding to antioxidant response elements (AREs) in the regulatory regions of its target genes (34). Our RNA-seq and proteomic data show increased expression of NRF2-regulated genes, including Heme Oxygenase 1 (HMOX1) and Glucose-6-phosphate dehydrogenase (G6PD), as well as SLC38A1, SLC38A2, and SLC38A3.

In addition to activating antioxidant genes to maintain redox balance, NRF2 acts as a metabolic switch by rerouting nutrient flux and driving metabolic reprogramming to enhance metabolic resilience, thereby increasing anabolic capacity and supporting rapid cell proliferation and survival under stressful conditions, including nutrient insufficiency (35, 36). STAT3 and HIF1α support NRF2-mediated metabolic reprogramming (37, 38). Our data demonstrate activation and nuclear localization of HIF1α and phosphorylated STAT3 (p-STAT3) in epithelial cells lining kidney cysts in TSC (**Fig. 4A-C**). NRF2 and STAT3 can interact to drive metabolic reprogramming and activate survival genes, including HIF1α (37–39). HIF1α can activate survival factors and glycolytic enzymes in proliferating cells (38, 39). Together, these molecules play a critical role in activating glutaminolysis and aerobic glycolysis, which are crucial for supplying nutrients to proliferating cystic epithelial cells and for cyst expansion in TSC.

**Figure 4.**
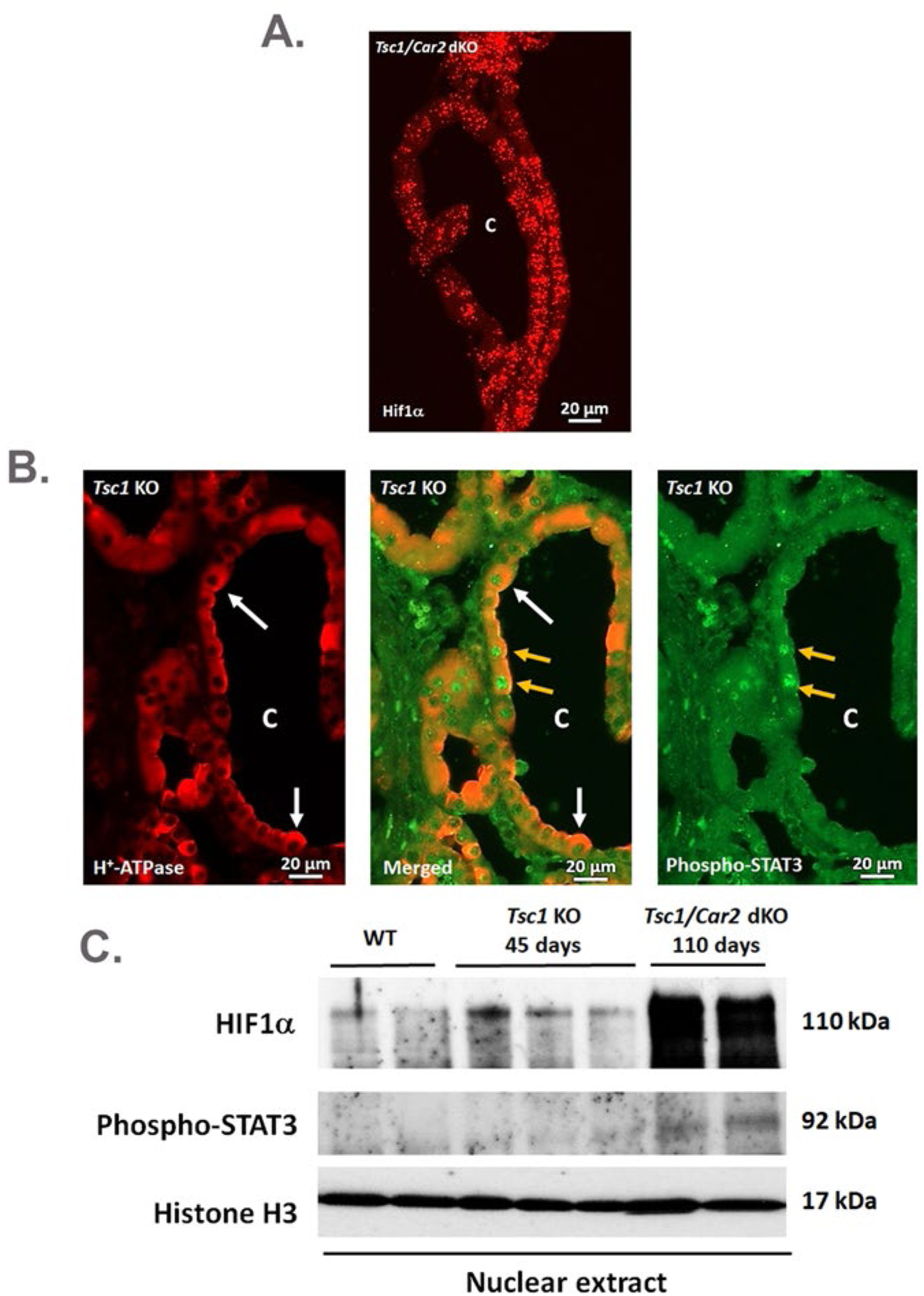
Localization and expression of HIF1α and phosphorylated-STAT3 in TSC mouse models. **A)** Fluorescence *in situ* hybridization (FISH) labeling of HIF1α lining the cyst epithelium in *Tsc1/Car2* dKO mice. “C” represents cysts. Scale bar = 20μm. **B)** Double immunofluorescence microscopy of H^+^- ATPase (red; white arrows) and phosphorylated-STAT3 (green; yellow arrows), with the merged image (middle) in *Tsc1*KO mice. “C” represents cysts. Scale bar = 20μm. **C)** Western blots of HIF1α and phosphorylated-STAT3 in WT, *Tsc1* KO, and *Tsc1/Car2* dKO nuclear extracts.

In support of increased glutaminolysis in kidney cystic epithelium and its crucial role in TSC cystogenesis, we found robust ectopic SLC38A3 (SNAT3) induction at the basolateral membrane of cyst-lining cells in TSC mice (*Tsc1* KO and *Tsc1/Car2* dKO) (**Fig. 5A-B**), and reduced kidney cyst burden in *Tsc1* KO mice on a glutamine-free diet (**Fig. 5C**).

**Figure 5.**
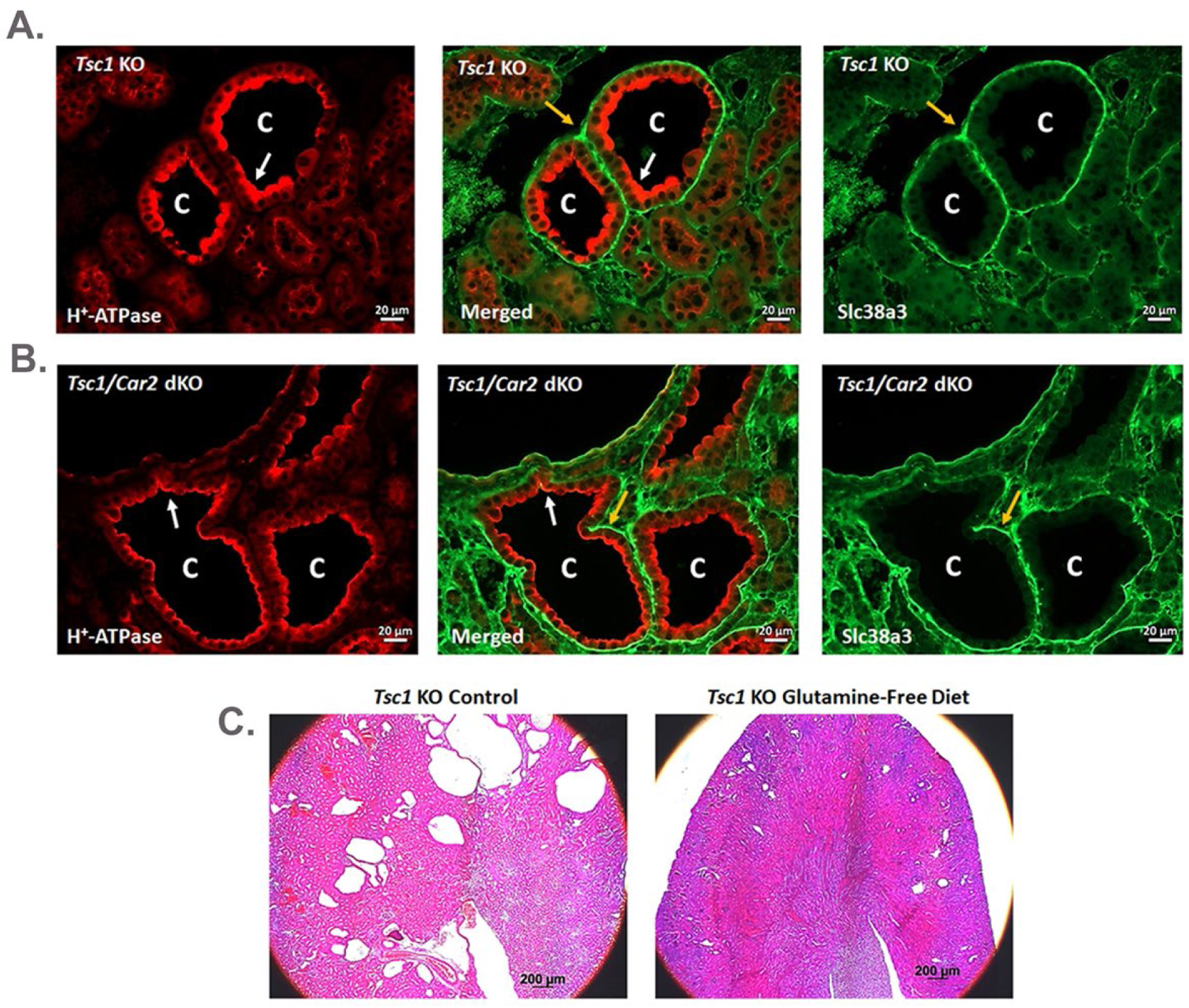
Glutaminolysis in TSC mouse models. Immunofluorescence localization of Slc38a3 (green; yellow arrows) and H^+^-ATPase (red; white arrows) in *Tsc1*KO **(A)** and *Tsc1/Car2* dKO **(B)** mice. Merged images are in the middle panel. “C” represents cysts. Scale bar = 20μm. **C)** Low magnification (4X) H&E images in *Tsc1* KO mice on regular diet (**left panel**) vs. *Tsc1* KO on a glutamine-free diet (**right panel).**

Our RNA-seq analysis and northern hybridizations showed upregulation of Rh Family Glycoprotein (RHBG and RHCG), the well-known NH_3_/NH_4_^+^ ammonia transporters/channels, on the membranes of A-IC cells lining the cysts (**Fig. 6A** and **6B**). Fluorescence in situ hybridization (FISH) showed a strong localization of RHCG to the cystic epithelial cells in *Tsc1* KO (**Fig. 6C**) and *Tsc1/Car2* dKO (**Fig. 6D**). Twenty-four-hour urine NH_3_/NH_4_^+^ excretion rates increased significantly in *Tsc1* KO vs. WT mice (37.29 ± 2.33 mmole in *Tsc1*KO vs 27.3 ± 2.23 mmole in WT mice; n=4; p<0.01).

**Figure 6.**
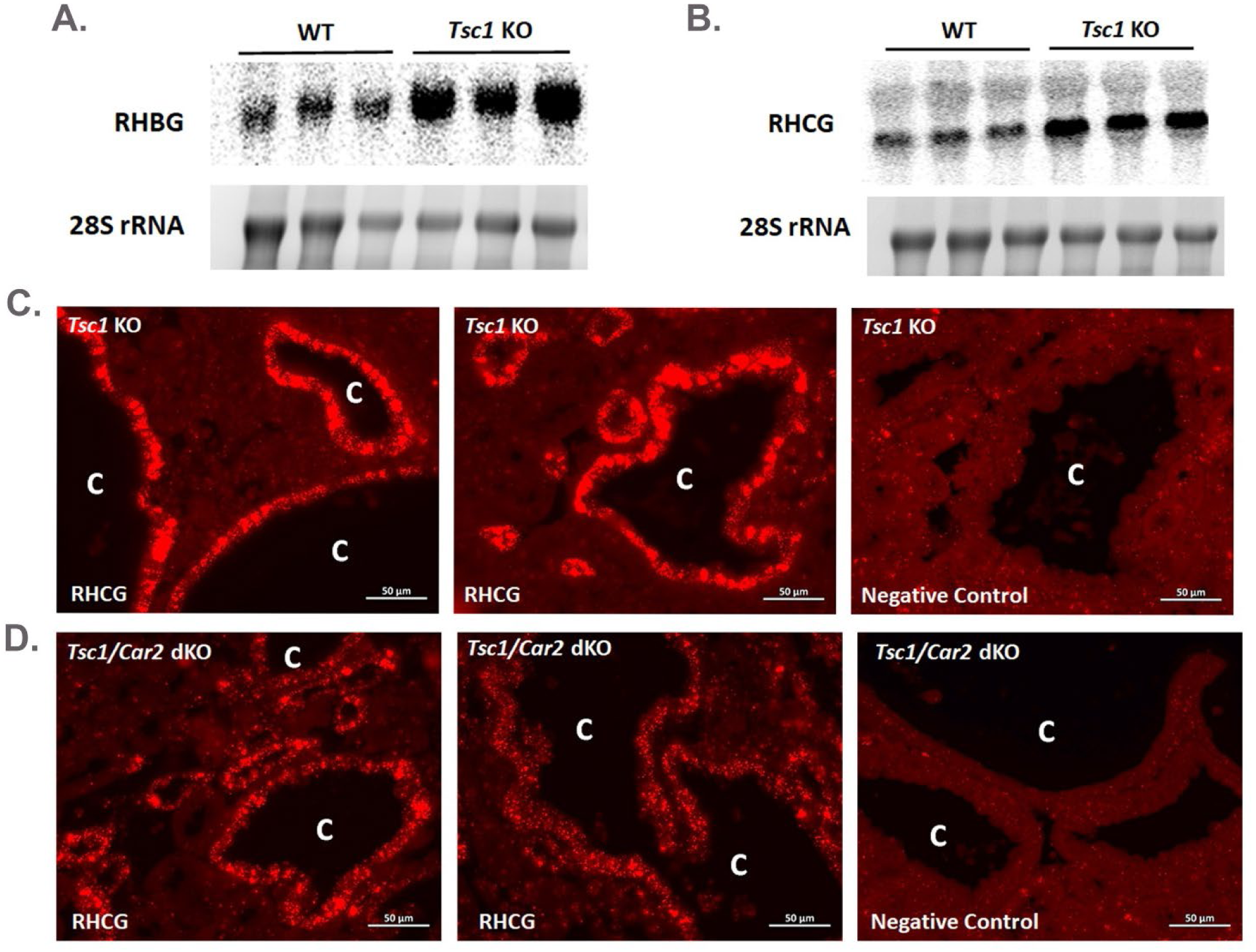
RHCG expression in TSC mouse models. Northern blots illustrating enhanced expression of RHBG **(A)** and RHCG **(B)** in kidneys of *Tsc1*KO compared to WT mice. Fluorescence *in situ* hybridization (FISH) of RHCG in cystic epithelial cells in **(C)** *Tsc1* KO (upper left and middle panels) and **(D)** *Tsc1/Car2* dKO (lower left and middle panels). Right panels in **C** and **D** are negative control images. “C” represents cysts. Scale bar = 50μm.

In addition to ectopic localization of SLC38A3, our RNA-seq studies as well as targeted RNA-scope experiments showed upregulation of several glutamine transporters, including SLC38A1 (SNAT1), SLC38A2 (SNAT2), SLC7A5 (LAT1), and SLC1A5 (ASCT2), in A-IC cells lining cysts in *Tsc1* KO and *Tsc1/Car2* dKO mice. Our RNA-seq analysis indicate upregulation of several aerobic glycolysis pathway molecules, including HK2, PFKFB3, and LDHA.

## Discussion

The trimolecular TSC1/TSC2/TBCD17 complex inhibits mTORC1 by inhibiting RHEB-GTPase (23–24). Mutations in TSC1 or TSC2 lead to the loss of RHEB-GTPase regulation, resulting in uncontrolled activation of mTORC1 (1, 2, 40, 41). Both ADPKD and TSC kidney cysts display mTORC1 activation (9, 12, 18). In TSC patients, inhibition of mTORC1 profoundly, but transiently, blunts the growth of kidney tumors and cysts (3, 9). Inhibition of mTORC1 in ADPKD patients, however, did not significantly improve kidney function or cyst volume (42, 43). Coupled with the contrasting cellular phenotype lining the cysts in TSC vs. ADPKD (12–20), these results highlight the distinct cystogenesis mechanisms in these two genetic kidney cystic disorders. Renal cyst epithelial cells in TSC mouse models and in humans with TSC display the loss of heterozygosity in only a small number of cystic cells (44–46).

### Molecules and pathways promoting kidney cystogenesis in TSC

The factors promoting kidney cyst generation and tumors in TSC are poorly understood. In addition to distinct differences in cells lining the cysts in TSC vs. ADPKD (12, 13, 18–20), expression studies demonstrate robust activation of the FOXI1 transcription factor and its downstream targets in the cyst epithelia of *Tsc1* (or *Tsc2*) KO mice, but not in *Pkd1*KO mice (12). FOXI1 is a master regulator of acid-base transporters in kidney intercalated cells, and its inactivation abrogated the cyst burden in the kidneys of *Tsc1*KO mice (12). The receptor tyrosine kinase c-KIT is activated in cystic epithelial cells of TSC mice in response to FOXI1 activation and is critical for phosphorylation-mediated inactivation of TSC2 and for promoting cell growth in TSC mouse kidneys (19, 20). Deletion of c-KIT in *Tsc1*KO mice abrogated kidney cystogenesis, and treatment with Imatinib, a c-KIT inhibitor, prevented cystogenesis in *Tsc1*KO mice (19, 20).

TFEB inactivation in the kidney distal nephron abrogates kidney cysts in *Tsc1/Ksp*-Cre mice (47). TSC2 regulates lysosome biogenesis via a TFEB-dependent mechanism (48). Deletion of *RagA/B* in kidney tubular cells triggers renal cystogenesis in TSC despite inhibiting mTORC1, consistent with TFEB important role in cystogenesis (49).

### The roles of antioxidant defense and metabolic reprogramming in kidney cystogenesis, cyst growth, and fluid secretion into the cyst lumen

mTORC1 activation increases metabolic demand, leading to oxidative stress and excess ROS generation (21, 22, 50–52). The transcription factor NRF2, as a master regulator of cellular redox homeostasis, controls antioxidant and anti-inflammatory defenses that protect renal cells against various injuries (24–26, 27–31). Under normal conditions, NRF2 is bound by KEAP1 in the cytoplasm and is targeted for destruction (23–31). However, when exposed to oxidative or electrophilic stress, specific cysteine sensors on KEAP1 are modified, resulting in its inactivation, which allows NRF2 to escape degradation and move to the nucleus, where it enhances the expression of cytoprotective and antioxidant genes; thereby, neutralizing the stress and restoring cellular balance (23–28). Mechanisms that can activate the NRF2 signaling pathway in proliferating cancer cells include: KEAP1 somatic mutations, epigenetic silencing of KEAP1, accumulation of P62, and transcriptional induction of NRF2 by oncogenic K-Ras and c-Myc (31–33). A critical pathway facilitating NRF2 rescue from KEAP1-mediated degradation is the **downregulation of Fumarate Hydratase 1 (FH1)** due to mTORC1 hyperactivation (30–33). This process can result in the accumulation of fumarate, which modifies **cysteine residues on KEAP1 via succination**; thereby, inactivating KEAP1 while activating NRF2 (28–32).

mTORC1 activation downregulates FH1 in specific tissues, including the renal epithelium (28–31), impairing the TCA cycle and causing accumulation of the oncometabolite fumarate (29–31), which drives epithelial transformation. The mechanistic connection between these two factors operates through a feed-forward oncogenic loop, whereby chronic upregulation of mTORC1 (as in TSC animal models or cells) downregulates FH1, sparing other TCA cycle enzymes (such as citrate synthase) (30, 31). Fumarate accumulation stabilizes HIF-1α and enhances cell proliferation (28, 29).

Our imaging studies in Figures 1 and 2 demonstrate robust and widespread nuclear NRF2 expression in kidney cyst epithelia in *Tsc1* and *Tsc2* KO mice, but not in *Pkd1* KO mice. Our results further indicate FH1 downregulation by proteomics and Western blot analysis, and KEAP1 succination in immunoprecipitation experiments in kidney lysates from mice with moderate or heavy cyst burden (Fig. 3A-C). These results are consistent with KEAP1 inactivation, which in turn stabilizes NRF2, promotes its nuclear localization, and ultimately activates antioxidant molecules, including HMOX1 and SLC38A1, SLC38A2, and SLC38A3 (Results).

### NRF2 expression in cystic epithelial cells in ADPKD

Few studies have examined the effect of NRF2 on kidney cystogenesis in ADPKD or TSC. A study in *Pkd1* (RC/RC) mice found that NRF2 localization in cystic epithelium varies with disease stage, with NRF2 showing nuclear localization around 30 days of age, followed by a significant decline at 180 days of age (53). The authors concluded that pharmacological agents that stabilize or induce NRF2 trigger antioxidant defense and slow cyst growth in *Pkd1* mutants (53).

### NRF2 expression in TSC cells

A study examining NRF2 expression demonstrated its upregulation and nuclear localization in TSC2-null cells (54). The authors used MEF (murine embryonic fibroblast) cells genetically engineered to exhibit TSC2 loss (55). These studies indicated robust nuclear localization of NRF2 along with markedly increased glutathione levels (54). While concerns have been raised about CDK7 inhibition in these studies (54), no concerns have been raised about robust NRF2 expression in TSC-deficient cells. In a renal cell cancer model generated by crossing a *Tsc*^lox/lox^ mouse with a Ksp-Cre mouse, the authors showed that mTORC1 activation could lead to FH1 downregulation and eventual NRF2 activation (28–31). Recent studies demonstrate NRF2 activation and antioxidant defense in TSC2-deficient cells, indicating NRF2’s protective role against cell death (56).

### Metabolic reprogramming

In addition to activating antioxidant genes to maintain redox balance, NRF2 acts as a metabolic switch by rerouting nutrient flux and driving metabolic reprogramming; thereby, supporting rapid cell proliferation under nutrient insufficiency (35, 57–59). To that end, NRF2, along with STAT3 and HIF1α, is activated in many cancers, where they promote glutaminolysis and aerobic glycolysis (59, 60). Our data demonstrate activation and nuclear localization of p-STAT3 and HIF1α in A-IC cells lining kidney cysts in TSC mouse models (Fig. 4A-C). NRF2 and STAT3 or NRF2 and HIF1α can interact to drive metabolic reprogramming and the activation of survival genes in proliferating cells. Together, these molecules can play a critical role in activating glutaminolysis and aerobic glycolysis, which are crucial for supplying nutrients to proliferating cystic epithelial cells and for cyst expansion in TSC.

Recent studies have explored the role of glutaminolysis in cell growth in cancerous tissues, particularly under catabolic stress conditions such as hypoxia or injury (58–60). Glutamine transporters, including SLC38A1 (SNAT1), SLC38A2 (SNAT2), SLC7A5 (LAT1), and SLC1A5 (ASCT2), are upregulated in proliferating kidney cystic epithelial cells in TSC (Results), consistent with their established role to fuel glutaminolysis (which sustains rapid growth), macromolecule synthesis, and mTOR activation (59–64). These transporters are investigated as targets for cancer chemotherapy. Few epithelial tumors show SLC38A3 activation in proliferating cells (59).

### The NRF2 addiction phenomenon

The KEAP1-NRF2 system is a pivotal defense mechanism against oxidative and electrophilic stress. Because NRF2 activation increases the antioxidant capabilities of cancer cells, persistently high NRF2 activity enhances tumor proliferation. NRF2 also drives metabolic reprogramming, establishing cellular metabolic processes that promote cell proliferation in concert with other oncogenic pathways. As a result, cancer cells with persistent NRF2 activation often develop “NRF2 addiction” (60, 61). Inhibition of NRF2 is a promising therapeutic approach for NRF2-addicted cancers.

### The Glutamine Addiction

Because proliferating cells constantly use glutamine to supply vital carbon and nitrogen for rapid DNA/RNA/protein synthesis, they become heavily dependent on an external glutamine supply **(glutamine addiction)** to replenish glutamine and generate fuel/nutrients (62, 63). The ectopic induction of SLC38A3 on the basolateral membrane of A-IC cells lining the kidney cysts in TSC mice points to a unique role for SLC38A3 as well as other glutamine transporters in glutaminolysis, enhanced cell proliferation and cystogenesis (Results; Fig. 5).

### Are the NRF2 Addiction and Glutamine Addiction Connected?

While they refer to two distinct states, NRF2 addiction and glutamine addiction are deeply intertwined, with NRF2 addiction directly driving glutamine addiction by creating a metabolic bottleneck that forces proliferating cells to consume glutamine just to meet NRF2’s antioxidant demands. Hyperactive NRF2 forces the cell to import glutamine, generate glutamate and large amounts of the antioxidant glutathione, and produce abundant citrate.

The co-localization of NRF2 and SLC38A3 in cystic epithelial cells in TSC (Fig. 5) and the known role of NRF2 in driving SLC38A3 expression by binding to the ARE in its promoter under oxidative stress (64, 65) point to a unique role for SLC38A3-mediated glutamine import in glutathione production and providing fuel for the TCA cycle, resulting in increased proliferation of A-IC cells lining the cysts. Coupled with the upregulation of the NH_3_ transporters **RHCG and RHBG** in (Fig. 6A-D) and GLS, these results point to reconstitution of the glutamine metabolic pathway in cystic epithelial cells; starting from glutamine influx (via SLC38A3 in Fig. 5, as well as other glutamine transporters; see above) to conversion to glutamate (and α-Ketoglutarate) in the mitochondria (via glutaminase; GLS) to NH_3_/NH_4_ ^+^ generation and export into the cyst lumen (**Schematic Diagram** in **Fig. 7**). NH*_3_* secretion across the apical membrane of A-IC cells requires activation of H^+^-ATPase to trap NH_3_ in the lumen by forming NH_4_ ^+^ (66; **Fig. 7**). It is very plausible that NH_3_ export into the cyst lumen and its trapping by H^+^-ATPase are critical rate-limiting (bottleneck) steps required for NH_3_ elimination and intra/extracellular pH regulation (**Fig. 7**); thereby, facilitating the continuation of glutamine import and metabolism, ultimately increasing the proliferation of A-IC cells lining the cysts.

**Figure 7.**
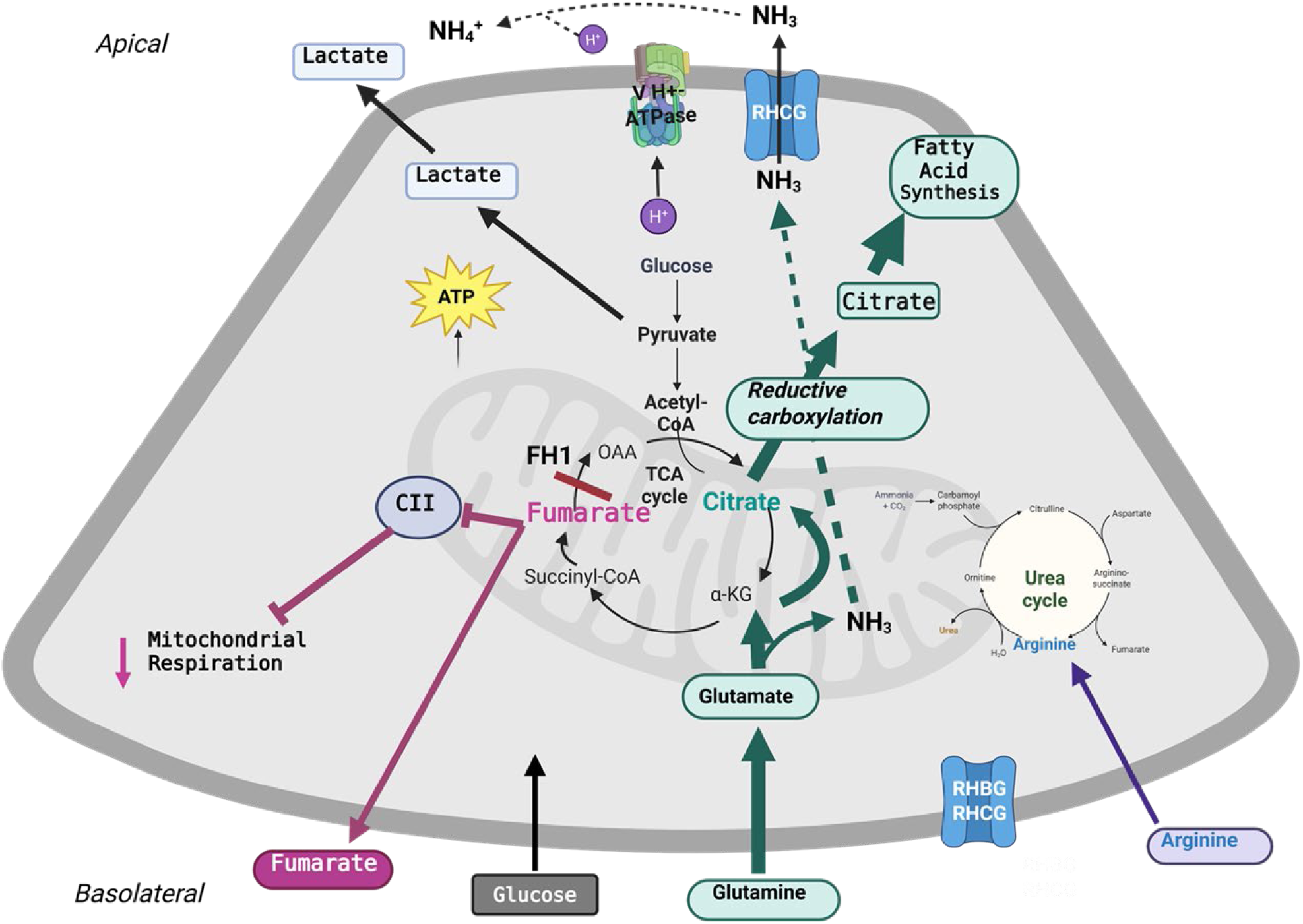
Schematic diagram depicting the reconstitution of the glutamine metabolic pathway in cystic epithelial cells. This pathway includes the import of glutamine which is converted to glutamate and NH_3_ generation, eventually leading to NH3 secretion across the apical membrane via RHCG in A-IC cells ling cysts in TSC mice.

In conclusion, our studies support a critical role for the canonical KEAP-NRF2 axis in kidney cystogenesis and point to an important role for FH1 downregulation, which increases fumarate and promotes KEAP1 succination, leading to KEAP1 inactivation and NRF2 nuclear localization. NRF2, in collaboration with STAT3 and HIF1α, is crucial in metabolic reprogramming. The ectopic induction of SLC38A3 in cystic epithelial cells, along with upregulation of RHBG and RHCG, and blunted cystogenesis in response to a glutamine-free diet point to the crucial role of glutaminolysis in kidney cystogenesis in TSC.

## Materials and Methods

### Animals

All animal experiments in this manuscript were conducted according to the “Guide for the Care and Use of Laboratory Animals” under protocols approved by the University of New Mexico Institutional Care and Use Committee (IACUC; 26-201742-HSC and 25-201620-HSC). Mice were group-housed in rooms with a controlled temperature of 22°C and with a 12hr light/12hr dark light cycle. IACUC approved training was received by all animal handlers. Mice were euthanized with an excess of pentobarbital sodium according to institutional guidelines and approved protocols.

### Generation of Tsc mouse models

Multiple TSC mouse models have been utilized in these studies (Table 1). These include: *Tsc1^f/f^* (*Tsc1* KO), *Tsc1/Car2* dKO, and *Tsc2^f/f^* (*Tsc2* KO). *Tsc1* KO mice (Jackson Labs, #005680; Bar Harbor, ME) were generated by crossing *Tsc1^f/f^* mice with *Aqp2* Cre mice (Jackson Labs, #006881) to eliminate *Tsc1* in kidney principal cells (67; 68). To generate the *Tsc1/Car2* dKO mice, *Tsc1* KO mice were crossed with *Car2* global KO mice. Similar to *Tsc1* KO, *Tsc2* KO mice were produced by mating *Tsc2^f/f^* (Jackson Labs, #027458) mice with *Aqp2* Cre mice; thereby, eradicating *Tsc2* in kidney principal cells (69). C57Bl6 mice (WT; Jackson Labs #000664) were used as controls.

### Genotyping

DNA was isolated from tail clippings of mice for PCR genotyping. Genotyping for *Tsc1^f/f^*, *Tsc2^f/f^*, as well as *Aqp2* Cre mice have previously been described (12, 30, 68–69).

### Tissue Collection

Mice were euthanized with an overdose of pentobarbital sodium according to our approved protocol. One kidney was quickly removed, snap frozen in liquid nitrogen and stored at −80° C for later protein extraction. The other kidney was placed in 4% paraformaldehyde in phosphate-buffered saline (PBS) overnight at 4°C then transferred to 70% ethanol for 24hrs before paraffin-embedding and sectioning to be used for microscopy analyses.

### RNA Seq analysis

RNA-seq and bioinformatic analyses were performed by Novogene Bioinformatics Technology Co., Ltd. (Sacramento, CA, USA). Briefly, total RNA was isolated from the kidneys and subjected to quality control analysis, using an Agilent 2100 Bioanalyzer with RNA 6000 Nano Kits (Agilent; Colorado Springs, CO, USA). After poly A selection, the samples were fragmented and reverse-transcribed to generate complementary DNA for sequencing. Libraries were sequenced on the HiSeqTM 2500 system (Illumina; San Diego, CA, USA). Clean reads were aligned with a mouse reference genome using Hisat2 v2.0.4. Gene expression levels were estimated using fragments per kilobase of transcript per million mapped fragments (FPKM) using HTSeq v0.9.1. 4.8.

### Proteomic analysis

The kidney proteomes of WT, *Tsc1*KO and *Tsc1*/*Car2*dKO were analyzed by the UNM Proteomic Core Facility using a SCIEX Triple Quad™ 5500+ LC-MS/MS instrument. The comparison of renal proteomes was performed using the FragPipe online tool (http://fragpipe-analyst.nesvilab.org/; accessed on 25 June 2026).

### Western Blot

Proteins were isolated in a combination of 10mM Triethaolamine and 250mM sucrose solution adjusted to pH 7.6 containing HALT protease and phosphatase inhibitor (ThermoFisher Scientific, Waltham, MA) and spun at 4,000g at 4°C for 10 minutes. The supernatant was removed and protein concentrations were measured with a BCA Protein Assay Kit (ThermoFisher Scientific). Proteins were loaded onto a 4-12% Novex Tris Glycine gel (ThermoFisher Scientific) and run at 100 volts and later transferred to a nitrocellulose membrane (ThermoFisher Scientific) at 25 volts for 90 minutes. Antibodies were diluted in a 5% milk blotting solution, added to the membrane, and allowed to incubate overnight at 4°C. Secondary antibodies were added the following day, washed, and developed with SuperSignal West Pico Plus Chemiluminescence substrate (ThermoFisher Scientific). Images were acquired using autoradiography film (Alkali Scientific; Ft. Lauderdale, FL).

### Immunofluorescence and Immunohistochemical Microscopy

Slides were baked at 60°C for 2 hours, allowed to cool at room temperature then deparaffinized in xylene. Next, the slides were rehydrated by placing them in decreasing percentages of ethanol, and they underwent an antigen retrieval protocol utilizing 10mM of sodium citrate and a short incubation in 0.85% NaCl. Slides were washed multiple times in PBS and incubated overnight in a humidity chamber at 4°C.

For immunofluorescence, slides were washed in PBS and incubated in secondary antibodies (Alexa Fluor 594 goat-anti-rabbit #A11037 1:200 and Alex Fluor 488 goat-anti-mouse #A11029 1:200; ThermoFisher Scientific) for 2 hours at room temperature. They were allowed to dry, cover-slipped with Vectashield Hardset mounting media (Vector labs #H-1500; Newark, CA), and images were obtained using a Zeiss LSM800 Airyscan microscope with Zeiss Zen software (Version 2.6).

For immunohistochemistry, slides were washed in PBS and treated with Vectastain Elite ABC Kit (Vector Labs, #PK-6101) and later stained with Vector VIP Substrate Kit (Vector Labs, #SK-4600) according to the manufacturer’s instructions. Slides were dried and cover slipped with VectaMount Permanent Mounting Medium (Vector Labs, #H-5000). Images were obtained with a Olympus CellSans Apex microscope with CellSans software.

### Fluorescence in situ Hybridization (FISH)

Tissue was paraffin-embedded and samples were prepared for labeling according to the manufacturer’s protocol (Advanced Cell Diagnostics, Newark, CA). Slides were hybridized with RNA-scope Multiplex Fluorescence V2 probes: Mm-Hif1α (Advanced Cell Diagnostics; #313821) and Mm-Rhcg (Advanced Cell Diagnostics; #1240961). TSA Vivid fluorophore 570 (Advanced Cell Diagnostics, #323272; 1:1000) was used for visualization. Slides were air-dried and cover-slipped using Vectashield HardSet mounting media (Vector Laboratories; #H-1500). Confocal imaging was carried out in the University of New Mexico Cancer Center Fluorescence Microscopy Shared Resource Center using an LSM800 Airy-Scan confocal microscope (Zeiss; Dublin, CA) and Zeiss Zen software (Version 2.6).

### Data Availability

The datasets and the data presented in this manuscript are available from the corresponding author upon request. Datasets utilized and/or analyzed in the current study are described in this manuscript. The RNA-SEQ data sets (Datasets EV1-2) have been submitted to the NCBI-Gene expression omnibus (GEO) repository.

- **DataSet EV1.** 1) RNASeq for Tsc1 KO vs WT Day 28; 2) GEO database (Accession number GSE311308)
- **DataSet EV2.** 1) RNASeq for Tsc1 KO vs WT Day 45; 2) GEO database (Accession number GSE311309) ***Statistical Analysis.*** The statistical differences between mean values +/− SD of multiple samples were determined using a one-tailed unpaired Student’s *t*-test. A *p-*value of less than 0.05 was considered statistically significant.

## Acknowledgments.

These studies were supported by Merit Review Award 5I01BX001000 (Department of Veterans Administration) (M.S.), Dialysis Clinics Inc. Grant C-4149 (M.S.), Department of Defense grant: TS240031(M.S.), and NIH Grant: NHLBIT32HL007736 (T. Resta, PI; M.S.). M.S. is a Senior Clinician

Scientist Investigator with the Department of Veterans Health Administration. This research made use of the Fluorescence Microscopy and Cell Imaging Shared Resource, which is supported partially by the University of New Mexico (UNM) Comprehensive Cancer Center Support Grant NCIP30CA118100, and the Human Tissue Repository & Tissue Analysis Shared Resource, which is partially supported by the UNM Comprehensive Cancer Center Support Grant NCI P30CA118100. The funders had no role in study design, data collection, analysis, decision to publish, or preparation of this manuscript.

